# Genomic insights into the evolution of secondary metabolism of *Escovopsis* and its allies, specialized fungal symbionts of fungus-farming ants

**DOI:** 10.1101/2023.11.06.564974

**Authors:** Aileen Berasategui, Hassan Salem, Abraham G. Moller, Yuliana Christopher, Quimi Vidaurre-Montoya, Caitlin Conn, Timothy D. Read, Andre Rodrigues, Nadine Ziemert, Nicole Gerardo

## Abstract

The metabolic intimacy of symbiosis often demands the work of specialists. Natural products and defensive secondary metabolites can drive specificity by ensuring infection and propagation across host generations. But in contrast to bacteria, little is known about the diversity and distribution of natural product biosynthetic pathways among fungi and how they evolve to facilitate symbiosis and adaptation to their host environment. In this study, we define the secondary metabolism of *Escovopsis* and closely related genera, members of which are specialized, diverse ascomycete fungi best known as mycoparasites of the fungal cultivars grown by fungus-growing ants. We ask how the gain and loss of various biosynthetic pathways corresponds to divergent lifestyles. Long-read sequencing allowed us to define the chromosomal features of representative *Escovopsis* strains, revealing highly reduced genomes (21.4-38.3 Mb) composed of 7-8 chromosomes. *Escovopsis* genomes are highly co-linear, with genes localizing not only in the same chromosome, but also in the same order. Macrosynteny is high within *Escovopsis* clades, and decreases with increasing phylogenetic distance, while maintaining a high degree of mesosynteny. To explore the evolutionary history of biosynthetic pathways in this group of symbionts relative to their encoding lineages, we performed an ancestral state reconstruction analysis, which revealed that, while many secondary metabolites are shared with non-ant associated sordariomycetes, 56 pathways are unique to the symbiotic genera. Reflecting adaptation to diverging ant agricultural systems, we observe that the stepwise acquisition of these pathways mirrors the ecological radiations of attine ants and the dynamic recruitment and replacement of their fungal cultivars. As different clades encode characteristic combinations of biosynthetic gene clusters, these delineating profiles provide important insights into the possible mechanisms underlying specificity between these symbionts and their hosts. Collectively, our findings shed light on the evolutionary dynamic nature of secondary metabolism in *Escovopsis* and its allies, reflecting adaptation of the symbionts to an ancient agricultural system.

## 1. Introduction

All parasites are specialists. At broad scales, each parasite can exploit some host species and not others. At finer scales, many parasite strains may be specialized on particular host genotypes within a species (1). While parasite host range is constrained by different evolutionary processes, including tradeoffs and arms race coevolutionary dynamics (2, 3), the molecular mechanisms underlying pathogen specialization and the evolutionary ecology of specificity have yet to be clearly linked. Similarly, little is known about the genomic architecture underlying the evolution of parasite specialization, the genomic consequences of host shifts and the genetic basis of shifts along the parasitism to mutualism continuum that underlies most symbioses.

Fungal symbionts are genetically tractable models for the study of host fidelity due to their diverse lifestyles and the occurrence of very closely related species that differ from each other primarily in their host range (4). Secondary metabolites, small molecules that are not necessary for the growth of an organism but aid in survival, play essential roles during fungal infection (5) and are known to affect the niche breath of fungal pathogens (4, 6, 7). Typically, specialists harbor a contracted array of specialized metabolites relative to generalists (4), reflecting the metabolic constraints that they experience in attempting to exploit different hosts. Given their role in mediating species interactions, secondary metabolites are central to arms-races dynamics (8, 9). Thus, their origin and distribution can reflect adaptation to specific host environments (7).

*Escovopsis* (Hypocreales: Hypocreaceae) is a specialized (10–12), diverse group of fungi found in the gardens of fungus-farming ants (Hymenoptera: Attini) (13). Currently, there are nine described species (14), some of which have been well-studied for their ability to parasitize the ants’ fungal cultivars (10–12). *Escovopsis* sp. can be virulent parasites of fungus-growing ant agriculture, causing garden biomass loss and colony decline (13, 15, 16). While it is presumed that most species in the group are similarly virulent, infection by certain species appear to be not as lethal, suggesting that the ecological role and evolutionary implications of these symbionts are not fully understood (17–19). In recognition of their morphological and ecological diversity, recent work has proposed splitting the *Escovopsis* genus into multiple genera (i.e., *Escovopsis, Luteomyces and Sympodiorosea*) (14). Here, we refer to all members of the group with the common name escovopsis for simplicity.

Fungus-farming ants are a monophyletic group of obligate agriculturalists (20). Attines feed their cultivated fungi (“cultivars”) with vegetative material, and in turn the cultivar represents the ants’ primary food source. Different attine lineages practice different modes of agriculture, exhibiting a high degree of specificity towards their cultivars (21, 22), and these different agricultural systems are generally associated with different *Escovopsis, Sympodiorosea* and *Luteomyces* spp. (23). The ancestral system, *lower agriculture*, is practiced by a group of ants that cultivate fungi in the Agaricales family. While most of the ant species in this system grow their cultivars in the form of mycelium, some ants in the lower agriculture system subsist on Agaricales that grow in yeast form, giving rise to the name of *yeast agriculture*. While *Sympodiorosea* and *Luteomyces* infections of mycelial-growing lower attine ant gardens are common, infection of yeast gardens has never been found (20). The third agricultural system is known as *coral agriculture,* in which a group of ants within the *Apterostigma* genus exploits fungus in the *Pterulaceae* family. Infection of coral gardens are also common, and include infection by *Escovopsis*, *Luteomyces* and other related taxa (10, 14). While lower attines, practicing lower, yeast and coral agriculture, are characterized by providing their cultivars with dead vegetative material, higher attines (practicing generalized higher agriculture and leaf-cutter agriculture) provide their fungal mutualists with freshly cut vegetative material (20). The two agricultural systems of higher attines are characterized by the obligate lifestyle of the cultivar, which cannot survive without association with the ants. *Generalized higher agriculture* is practiced by ants cultivating a derived clade of agaricaceous fungi, whereas in the most derived agricultural system, that of *leaf-cutter agriculture*, a single fungal species *Leucoagaricus gongylophorus* is responsible for ant survival. Higher agriculture gardens are commonly infected with *Escovopsis*, most of which are *Escovopsis* spp. closely related to the best studied species, *E. weberi* (24).

Escovopsis symbionts show a high degree of host fidelity, being able to infect some cultivars but not others. This degree of partner specificity suggests a long history of coevolution, as demonstrated by the phylogenetic congruence between attines, their cultivars and *Escovopsis*, particularly at the broad interspecific scale (10). To manage infections, ants actively weed infected portions of garden, and many attine species associate with actinomycete *Pseudonocardia* that synthesize antifungal compounds that inhibit escovopsis growth (25).

Despite consistent patterns of co-diversification across the tripartite interaction between the ants, their cultivars and escovopsis (10), and the outsized role of natural products in mediating fungal specialization, the secondary metabolism of escovopsis remains relatively unexplored relative to the evolutionary ecology of attine ants and their cultivars. Only a few *Escovopsis*-derived compounds have been identified (26, 27), though recent genome annotation indicates the potential to produce many more (28). Here, we performed long-read genome sequencing, assembly, and annotation to describe the chromosomal architecture, conservation, and organization of escovopsis, which will facilitate future annotation of the biosynthetic machinery. After defining the secondary metabolism across the group and spanning representative host ranges, we contextualize the distribution of biosynthetic gene clusters relative to patterns of specialization and fidelity. Through comparative genomics, extensive manual curation of biosynthetic gene clusters, and ancestral state reconstruction, we outline a symbiont whose secondary metabolism broadly reflects the dynamic patterns of cultivar recruitment and replacement by attine ants.

## 2. Material and Methods

### a. Sample collection, isolation, DNA extraction and genome sequencing

Strains of escovopsis infecting all attine agricultural systems were obtained from the Emory collection (Table S1). To obtain DNA, fungi were grown on PDA plates at room temperature. Genomic DNA was extracted by crushing fungal tissue with liquid nitrogen and subsequently isolating the DNA using a phenol-chloroform protocol (29). Sequencing was performed on a HiSeq 2500 Sequencing system from Illumina, utilizing the paired-end 150 bp technology. Both library preparation and DNA sequencing were carried out at Novogene. Additionally, DNA from strains NGL095 *(E. weberi),* NGL070 *(E. multiformis)* and NGL057 (*Hypocreales: Hypocreaceae*, cf. *Escovopsis*) was also sequenced with PacBio technology by OmegaBioservices.

### b. Genome assembly and annotation

Strains sequenced with PacBio technology were assembled with Canu v.1.8 (30) and polished with their corresponding Illumina reads using Pilon v.1.23 (31). Those strains sequenced with Illumina alone were quality checked with FastQC (32), trimmed with trimmomatic (33), and subsequently assembled with Spades v.3.13.0 (34). Genome assembly quality was evaluated using BUSCO v.3 (35). GC content was calculated with the script GC_content.pl by Damien Richard (https://github.com/DamienFr/GC_content_in_sliding_window/ [last accessed July 2023]), using default parameters. The genomic dataset was completed with the addition of six previously sequenced *Escovopsis* genomes (26), as well as 14 closely related species from the Hypocreales family obtained from JGI Mycocosm (Table S1). The highly contiguous hybrid assemblies NGL070, NGL095 and NGL057 were screened for stretches of telomeric repeats (TTAGGG)n at the end of contigs, and contigs harboring these repeats at both ends were considered complete chromosomes.

To compare genomic architecture conservation between *Escovopsis* strains, a synteny analysis was performed on the proteome sets of the most unfragmented assemblies in our dataset employing GENESPACE v0.9.3 (36) as implemented in R. This dataset comprised the three hybrid assemblies NGL095, NGL070 and NGL057, as well as EACOL, EAECHC, EAECHR, ETCORN and EATTINE.

All assemblies were subjected to gene prediction and annotation using the Funannotate v.1.8.3 (37) pipeline. Repeats were identified with RepeatModeler and soft masked using RepeatMasker. Protein evidence from a UniprotKB/Swiss-Prot-curated database (38), and the proteomes from *Trichoderma sp., Cladobotryum sp., Hypomyces rosellus* and *H. perniciosus* were aligned to the genomes using TBlastN and Exonerate (39). Three gene prediction tools were used: AUGUSTUS v3.3.3(40), snap (41) and GlimmerHMM v3.0.4 (42). tRNAs were predicted with tRNAscan-SE (43). Consensus gene models were found with EvidenceModeler (44). Functional annotation was conducted using BlastP to search the Uniprot/SwissProt protein database. Protein families (Pfam) and Gene Ontology (GO) terms were assigned with InterProScan5 (45). Additional predictions were inferred by alignments to the eggnog orthology database (46), using emapper v3 (47). The secretome was predicted using Phobius v.1.01 (48), which identifies proteins carrying a signal peptide. Carbohydrate active enzymes were identified using HMMER v3.3 (49) and family specific HMM profiles of the dbCAN2 server (50). Proteases and protease inhibitors were predicted using the MEROPS database v (51), and biosynthetic gene clusters were annotated using fungiSMASH v6 (52) with relaxed parameters.

### c. Phylogenetic reconstruction

The phylogenetic relationship of *Escovopsis* spp. and their close relatives was reconstructed using the BUSCO_phylogenomics pipeline (53). In short, single copy orthologues for each genome were identified by running BUSCO v5 (35) with the Ascomycota_odb10 lineage database. This analysis identified 660 single-copy orthologs shared by all 34 strains in the dataset. Gene sequences were aligned with MUSCLE (54), and the alignment was trimmed with TrimAl (55). Output alignments were concatenated into a supermatrix. A maximum likelihood tree was built with IQ-TREE (56), allowing ModelFinder (57) to predict the best evolutionary model for partitioning the alignment. The resulting tree was rooted using *Trichoderma spp.* and visualized with iTol v6 (58).

In order to place escovopsis strains in a broader phylogenetic context, we performed a multi-locus analysis employing different molecular markers (i.e., ITS, TEF and LSU). Sequences were aligned in MAFFT v.7 (59) and a phylogenetic tree was reconstructed using maximum likelihood (ML) in RAxML (60). The GTR nucleotide substitution model was selected by independent runs in jModelTest2 (Darriba et al. 2012) using the Akaike Information Criterion (AIC) with 95% confidence intervals. 1000 independent trees and 1000 bootstrap replicates were performed and the final tree was edited in FigTree v.1.4 and Adobe Illustrator CC v.17.1.

#### Gene Cluster Family (GCF) identification

Biosynthetic gene clusters (BGCs) of all fungal strains were identified again using FungiSMASH 6.1 (52) with relaxed parameters, utilizing as input the GenBank files obtained after genome annotation. With the aid of cblaster v.1.3.12 (61), BGCs split onto different contigs, especially those located on contig edges, were manually assembled based on homology with other BGCs in the dataset. Likewise, fused BGCs were manually split into separate BGCs. The final BGC set was analyzed using BiG-SCAPE v1.0.1 (62) to identify homologous BGCs across all strains and to cluster related BGCs into gene cluster families (GCFs). BGCs from the MiBIG database 2.0 (63) were included in the analysis with the –mibig flag to identify already described BGCs. The scikit-learn package was downgraded to v.0.19.1, and the following parameters were enabled: –mix, --hybrids-off, and –include_singletons. The program was run in ‘glocal’ alignment mode with edge-length cutoffs from 0.1 to 0.9, with step increments of 0.1. After inspection, networks at thresholds 0.5-0.6 were found to be similar and further analyses were based on a cutoff of 0.5. The resulting sequence similarity matrixes were visualized using Cytoscape v.3.9.0 (64). A presence/absence matrix was built to evaluate BGC distribution, with 1 representing presence and 0 absence of a GCF in a fungal strain and was visualized as a heatmap employing R. To assess whether BGC profiles can delineate groups of escovopsis, a Jaccard distance matrix was computed using the presence/absence table. The distance matrix was then used to construct nonmetric multidimensional scaling (NMDS) ordination plots to detect grouping patterns and subjected to a PERMANOVA analysis (permutational multivariate analysis of variance) to identify significant factors underlying observed groupings. To assess the adequacy of our sampling, and to provide an estimate of GCF richness for the given sequencing effort, rarefaction curves were built at the genus level, and at both levels of attine agricultural systems (*i.e.,* lower and higher agriculture, as well as lower, coral, general higher and leaf-cutter agriculture).

### d. Co-cladogenesis analyses

To assess whether BGC profiles delineate escovopsis, the GCF presence/absence was subjected to a hierarchical clustering analysis using a correlation-centered similarity metric with the complete linkage clustering method. A tanglegram was built in R to evaluate the congruency between the symbiont phylogeny and strain BGC profiles using the package “dendextend” v 1.17.1.

### e. Ancestral State Reconstruction

To assess the evolutionary history of the GCFs, the ancestral node for each GCF was inferred in the species tree using the trace character history function implemented in Mesquite (65). In some cases, BiG-SCAPE split BGCs into multiple GCFs that were highly homologous, suggesting they may be involved in the biosynthesis of related compounds. Data exploration with different BiG-SCAPE similarity cutoffs did not resolve these relationships, prompting the manual grouping of GCFs into pathways (66, 67). GCFs were considered to belong to the same pathway if: (i) the BGCs shared similar architecture, (ii) the majority of the genes in the cluster had the same function, albeit not necessarily in the same order, and (iii) the majority of genes in the BGC had a BLAST similarity of more than 50% over 80% coverage rate (67). A pathway presence/absence table was used as a character matrix, and likelihood calculations were performed using the Mk1 model. Likelihood scores >50% were used to infer the points of pathway acquisition in the species tree.

## 3. Results and Discussion

To characterize the genomic features and secondary metabolism potential of this diverse group of specialized symbionts, we sequenced the genomes of 14 strains across the symbiont phylogeny, spanning all ant-agriculture ecologies (Table S1) (20), with the exception of yeast agriculture where escovopsis have never been found. Three strains belonging to different clades were sequenced with PacBio and Illumina technologies, whereas the rest were sequenced with Illumina alone (Table S2). We expanded our dataset with the addition of 24 additional escovopsis strains that were publicly available) (26, 28), and a number of other closely related fungal species from the Hypocreaceae family (Table S1).

The quality of the genomic assemblies generated in this study was high, with an average BUSCO (Benchmarking Universal Single-Copy Orthologs) score of 94.7% for the Ascomycota lineage dataset (Table S2). GC content ranged from 47.2% to 56.4%, with an average of 52.3% (Table S2), consistent with recent reports (24, 28) and other Pezizomycotina fungi (68).

### Phylogenetics of *Escovopsis* and relatives

To infer a genome-scale phylogeny of *Escovopsis* and relatives, we employed a concatenation approach using single-copy genes. The inferred proteomes of all 52 species in our dataset were subjected to an orthology analysis, resulting in 2314 single-copy orthologous genes that were subsequently utilized to infer a phylogeny. Our data reveals that the attine-associated symbionts form a monophyletic group, sister to a clade composed of *Cladobotryum* sp. and *Hypomyces rosellus*, both mycoparasites (Figure 1A). The evolutionary history of escovopsis suggested by this phylogeny generally reflects that of the fungus-growing ants (20). As such, strains infecting gardens of lower attines appear basal to the rest, whereas most recently diverging lineages are associated with higher attine agriculture and leaf-cutter ants (Figure 1A). The shift experienced by some lower attines to cultivating Pterulaceae fungi is also mirrored by the phylogeny, with an intermediate clade exploiting coral agriculture, represented by strains NGL070, ICBG726, ICBG1054, ICBG1065 and ICBG1075. Highlighting the diversity of escovopsis associated with coral agriculture, a clade including four strains associated with coral fungi (ICBG712, ICBG721, NGL057 and NGL216), appear within the basal members of this monophyletic group (Figure 1A). The presence of these two distinct coral agriculture-associated clades, therefore, break congruence of the ant and escovopsis phylogenies. Recent studies have proposed to split the genus *Escovopsis* intro three different genera (*Escovopsis*, *Sympodiorosea*, and *Luteomyces*) (14). To assess whether these two coral agriculture-associated clades may in fact represent two distinct taxonomical genera, we inferred the phylogenetic position of the escovopsis in this study among those from previous studies (14). Our results (Figure S1) suggest that strains within these two clades indeed belong to different genera. Together with strains exploiting higher agriculture (*E. weberi, E. moelleri* and *E. aspergilloides*), the intermediate clade exploiting coral agriculture are true *Escovopsis* (*E. multiformis*). However, its sister clade contains strains closely related to the newly described *Sympodiorosea*. Interestingly, the basal-most clade comprises strains most closely related to *Luteomyces* and to strains belonging to an as of yet undescribed genus (Figure S1). Overall, these results highlight the need for further work to fully resolve the taxonomical diversity withing these group of parasites.

**Figure 1.**
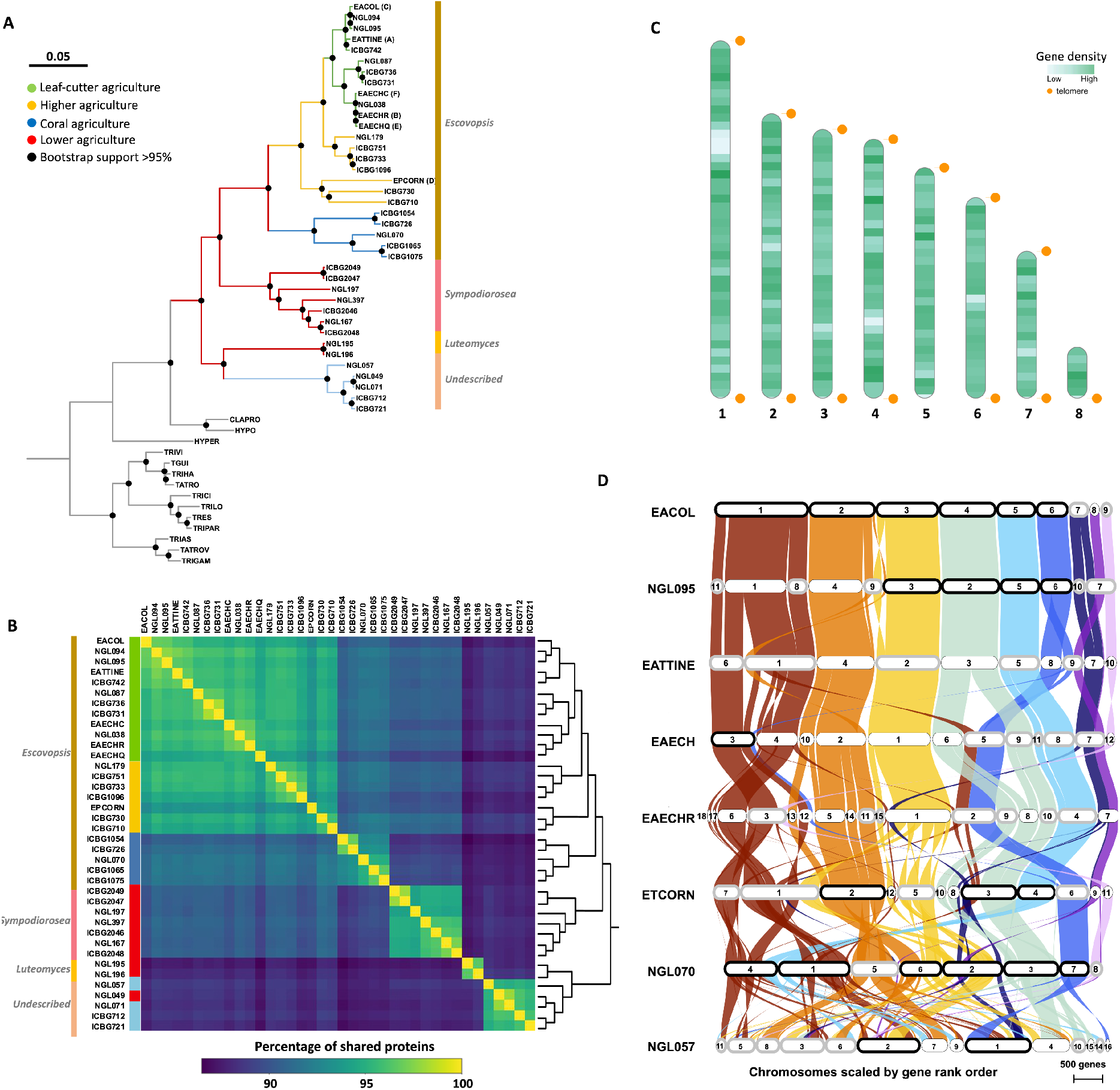
Genomic features of escovopsis. A) Phylogenomic tree of *Escovopsis* and allies constructed with a supermatrix approach on 2314 single-copy orthologous genes. Black dots represent bootstrap support higher than 90%. Branch colors describe different attine agricultural systems: green, leaf-cutter agriculture; yellow, general higher agriculture; blue, coral agriculture (divided in A, blue; and B, light blue; reflecting most derived and more basal coral agriculture); and red, lower agriculture. Side colored bars represent new taxonomical affiliations based on (14). (B) Heatmap depicting the percentage of conserved proteins across escovopsis strains. Lighter colors represent high levels of shared proteins, whereas dark colors depict fewer shared proteins. The dendrogram on the right represents a hierarchical clustering analysis. (C) Ideogram representing the chromosomal level assembly of an *Escovopsis* sp. strain isolated from an *Apterostigma dentigerum* nest (NGL070). Green colored bands represent regions with high gene density; lighter colors depict low gene density regions. Orange dots represent areas harboring telomeric repeats. (D) Synteny plot depicting the collinearity between different escovopsis genomes across attine agriculture. Highly syntenic regions are connected by colored bands. Contigs in black boxes represent complete chromosomes, whereas those in grey harbor telomeric repeats just at one chromosomal end.

To estimate the evolutionary distance between strains, we performed a POCP (Percentage of Conserved Proteins) analysis. As expected, with increased phylogenetic distance POCP values decrease. For instance, escovopsis infecting leaf-cutter agriculture share on average 96% of their proteins among each other, whereas only around 88% are shared with *Luteomyces*, *Sympodiorosea* and the newly undescribed genus (Table S3, Figure 1B). Despite appearing in the same clade in our phylogeny, *Luteomyces* strains and those from the undescribed genus, share as many proteins between each other (88%) as they do with strains parasitizing any other agricultural system. This suggests that there is as much phylogenetic divergence between these two groups, as there is between them and any other escovopsis clade, supporting the notion that what has been traditionally considered *Escovopsis* is in fact at least three, and possibly four, different genera. Furthermore, POCP values lower than 91% segregate our dataset into the recently proposed genera, whereas values above 91% and 95% delineate distinct species and strains within a species respectively (Table S3). Mirroring our phylogenetic placement of *M. zeteki*-associated *Escovopsis*, in POCP analysis, NGL179 shares more proteins (95.1%) with strains infecting leaf-cutter agriculture, than with those exploiting general higher agriculture (92%). POCP analyses have been useful to resolve bacterial groups at genus level, which correlate with POCP values < 50%. While some studies have implemented the method in some fungi at the family level (POCP values < 70%) (70), this strategy cannot be widely employed yet for delineating fungal groups, as genome sampling in fungi remains scarce. However, our POCP analysis reveals a significant degree of genetic diversity between *Escovopsis* (sensu lato) clades, and suggests a protein similarity threshold of 87-91% to delineate different genera in this group of parasites. Further efforts are required to elucidate whether the POCP differences can delineate distinct genera in a diversity of fungi.

### Escovopsis genomes are organized into highly syntenic chromosomes

To elucidate the genomic organization of escovopsis, we screened the genomes of the four most contiguous assemblies for telomeric repeats. In NGL070, stretches of (TTAGGG)n were detected at both ends of six contigs, representing complete chromosomes (Figure 1C). The two remaining contigs harbored telomeric repeats only at one end, constituting either two fragments of the same chromosome, or two distinct uncomplete chromosomes. A similar pattern was observed for the highly contiguous EACOL, NGL095 and NGL057 strains, harboring 6, 4 and 2 complete chromosomes, and 2, 5 and 7 fragmented ones with telomeric repeats at one end respectively (Figure 1D). These observations suggest that escovopsis has 7-8 chromosomes, in agreement with other members of the Hypocreales family, such as *Trichoderma reesei, Neurospora crassa* (71), and *Metarhizium brunneum* (72), which organize their genomes in seven chromosomes.

To assess the conservation of escovopsis genomic architecture, we performed a synteny analysis of the eight most continuous genomes available. Our ortholog-based analysis reveals that strains share a high degree of collinearity, with 87.83% of the genes appearing in the same chromosome and in the same order (Figure 1D). This is particularly apparent among strains of the same clade, as evidenced by those parasitizing leaf-cutter agriculture (EACOL, NGL095, EATTINE, EAECHC, and EAECHR). As expected, collinearity has a positive correlation with phylogenetic relatedness, with distant strains exhibiting increasingly different genomic organization. Chromosomes 1, 2, 3, 4 and 5 (nomenclature relative to strain EACOL), are extremely well conserved, extending beyond *Escovopsis* spp. infecting leaf cutter agriculture and including those involved in general higher agriculture. Chromosome 6, although well conserved in strains exploiting general higher agriculture and coral agriculture-associated NGL070, has experienced recent rearrangements, as evidenced by its fusion with a fragment of chromosome 1 occurring in the clade represented by EAECHC and EAECHR. Previous reports revealed a high degree of micro mesosynteny between genomes of *Escovopsis* and *Trichoderma* (24), suggesting that both genomes are organized in genome segments with similar gene content but rearranged in order and orientation.

### Escovopsis harbor a reduced genome

Fungi vary extensively in genome size, spanning three orders of magnitude and ranging from the small genomes of some Microsporidia (2Mb) to the large ones in Picciniales fungi (2Gb). Some of the smallest genomes are found in obligate parasites (73). Escovopsis genome sizes range between 21.4 and 38.3 Mbp (40.7Mb), with an average of 28.7 Mb, corroborating previous studies (24, 28) that estimated their genome sizes around 24.7-27.2 Mb. These genomes are reduced in size relative to those of closely related Sordariomycetes (Figure 2A, 2B, S2A, S2B). Interestingly, *Escovopsis* represents three of the five smallest genomes from all Sordariomycetes strains publicly available in Mycocosms (https://mycocosm.jgi.doe.gov) (Figure 2A). The other two belong to *Ophiocordyceps camponoti-rufipedis* and *O. australis* strain 1348a, both highly specific parasites of ants (74). Within escovopsis, lower attine *Luteomyces* spp. strains harbor significantly smaller genomes than those infecting higher attine nests (Figure 2B, Table S4A). No differences in genome size were detected in escovopsis exploiting other agricultural systems (Figure 2B, Table S4B), with the exception of strains infecting higher agriculture, which are highly variable.

**Figure 2.**
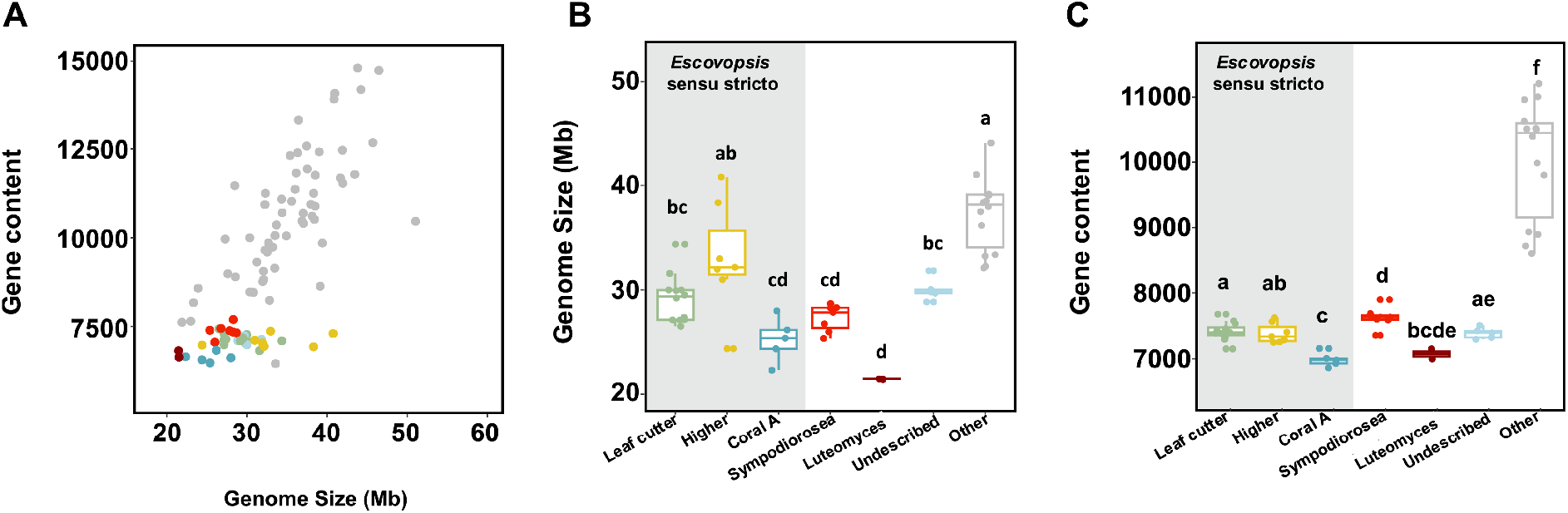
Escovopsis harbor reduced genomes with fewer genes than their non-ant associated relatives. (A) Relationship between genome size and gene content for sequenced fungal genomes. Genome size (B) and gene content (C) of escovopsis strains across different attine agricultural systems. Box colors denote attine clades systems: green, leaf-cutter agriculture (*Escovopsis* spp.); yellow, general higher agriculture (*Escovopsis* spp.); blue, coral agriculture A (*Escovopsis* spp.); red, lower agriculture (*Sympodiorosea* spp.); and light blue, coral agriculture B (undescribed genus), grey other *Sordariomycetes*.

Gene number in escovopsis ranged between 6477 and 7693 (Table S2), representing 9 out of the 10 species in Mycocosm with the fewest genes within the Sordariomycetes. Unlike other fungi in the family, where gene content positively correlates with genome size (r^2^ = 0.32, p < 0.0001), gene number in escovopsis is stable and does not associate with genome size (r^2^ = 0.06, p = 0.07) (Figure 2A). While escovopsis harbor fewer genes than their relatives (Kruskal-Wallis rank sum test, X^2^ = 30.11, d.f. = 1, p < 0.001, Figure S3A), there is no difference in gene content between escovopsis exploiting the nests of lower and higher attines (Table S4C, Figure S3B). However, those within the coral A clade have a slightly different gene content than those associated with lower and leaf-cutter agriculture (Figure 2C, Table S4D). These results are congruent with recent surveys (28) revealing that total coding sequence (CDS) length and intron number in escovopsis genomes are low relative to free-living relatives, consistent with reduced gene content.

In addition to gene number, we investigated two drivers of fungal genome size: repeat content and Repeat-Induced Point mutation. First, while transposable elements are often associated with fungal pathogens (75, 76), their number in escovopsis is significantly lower than in non-ant associated relatives (Kruskal-Wallis, X^2^ = 14.19, d.f. = 1, p < 0.001), which in part, explains escovopsis’ small genomes. Second, fungi have evolved a genome defense mechanism to mitigate the potentially detrimental consequences of transposable elements and other repeated genomic regions (77). By altering nucleotide ratios, Repeat-Induced Point mutation (RIP) leads to gene inactivation and genome reduction. Deactivation of RIP, therefore, can lead to genome expansion due to retrotransposon proliferation (78). Previous reports based on the analysis of just one strain, suggested that *E. weberi* may have lost genes involved in RIP. BLAST analyses with the sequences of the two canonical genes known to mediate the RIP pathway revealed that all *Escovopsis* genomes in our dataset harbored orthologs for one gene essential to the RIP process (RID, RIP deficient) but lacked orthologs to the other RIP canonical gene (DIM2, defective in methylation) (Table S5). Genome-wide RIP analyses using The RIPper’s sliding window approach revealed that all escovopsis strains show hallmarks of RIP (Table S6), although they vary greatly in the proportion of their genomes that are affected by it. While some strains harbored little evidence of RIP (ICBG1096, 1.01%), others are highly affected by it, with the most extreme case being ICBG1075, where 23.26% of its genome present hallmarks of RIP. This variation across escovopsis genomes of similar size indicates that RIP is not solely responsible for genome reduction in this group, but it may play some role in some species.

Escovopsis*’* small genomes and the genomic traces of RIP, together with the generalized loss of DIM2, support previous studies (24) that proposed RIP as a genomic defensive mechanism that limited transposon proliferation in *Escovopsis* in the past. A consequence of RIP is the relative absence of duplicated genes (77). Therefore, the loss of this defense mechanism, may represent an opportunity for this parasite to evolve new metabolic functions through gene duplication and subfuncionalization.

Ascomycota with genome sizes between 25 and 70 Mb, and in particular Sordariomycetes, often exhibit positive correlations between genome size and gene content (73, 79, 80). *Escovopsis* evades this trend (Figure 2A), suggesting that a different evolutionary processes may be affecting this genus. Symbiosis often leads to the streamlining of microbial genomes through genome reduction and gene loss, as epitomized by the tiny genomes of many bacterial endosymbionts of insects (81). Genome streamlining in bacteria can be explained by the loss of redundant genes with drift (82), or by selection against non-essential genes (83). Similar dynamics can occur in fungal mutualists and parasites (84, 85). In particular, fungal parasites associated with insects have been shown to be particularly prone to gene loss (80). Within the Sordariomycetes, the smallest genomes belong almost exclusively to endosymbionts, endoparasites, or fungal parasites vectored by insects (80).

### BGC Diversity and distribution

Secondary metabolites in fungi can define ecological niches (86), delimit host ranges (4, 87, 88), and provide selective advantages under specific ecological conditions (89). The metabolic pathways responsible for the synthesis of microbial toxins and other secondary metabolites are typically encoded by biosynthetic gene clusters (BGCs). BGCs encode for backbone enzymes responsible for the synthesis of the core structure of a metabolite, as well as tailoring enzymes that modify this assembly, along with transcription factors and transporters (90). To assess the biosynthetic potential of the escovopsis, we performed a computational genome mining analysis using the program fungiSMASH (52). The most common backbone enzymes in fungi include polyketide synthases (PKSs), nonribosomal peptide synthetases (NRPSs), terpene synthases, and dimethylallyltransferases (DMATs) (91). All escovopsis genomes analyzed harbored a diversity of BGCs belonging to the major biosynthetic classes (Table S7). Escovopsis*’* chemical potential content ranged from 16 BGCs in NGL197, to 33 in NGL057. On average, each genome featured 23 BGCs, and an average metabolic diversity of 28.7% NRPs, 25.6% PKS, 21.3% terpenoids, 16.3% hybrids, 2.4% betalactones and 3.6% others. There was no correlation between the number of BGCs and the number of contigs or scaffolds per genome (R^2^ = 0.03, p = 0.13), suggesting that our dataset was robust and that the different sequencing technologies employed did not bias our BGC survey. In addition, no correlation was found between the number of BGCs in each escovopsis strain and genome size (R^2^=0.004, p = 0.7).

While fungi within the Hypocreales are prolific secondary metabolite producers, with an average of 43 BGCs per genome, escovopsis have significantly fewer BGCs than their non-fungus-farming ant associated relatives (Kruskal-Wallis X^2^ = 28.17, d.f. = 1, p < 0.001, Figure S4A), corroborating recent findings using fewer escovopsis genomes (28). We found no statistical differences in BGC abundance between escovopsis strains infecting higher or lower attine nests (Figure S4B), nor between the majority of strains attacking different agricultural systems (Kruskal-Wallis p=0.67, Figure 3A), with the exception of small differences in BGC number in strains infecting general higher agriculture and leaf-cutter agriculture. Upon graphical inspection, we observed a clear bimodal distribution in BGC abundance in strains infecting lower agriculture (Figure 3A), that unequivocally divided the dataset in distinct phylogenetic taxa. We therefore explored whether there is a correlation between BGC content and phylogeny by assessing differences in BGC number across escovopsis groups (Figure S4C). All clades harbored significantly different number of BGCs, with the exception of *Leutomyces* spp. and the undescribed genera, both composed of strains infecting lower agriculture (Kruskal-Wallis, X^2^ = 47.91, d.f. = 6, p < 0.001). Strains within *Luteomyces* and the undescribed new genus (i.e., NGL195, NGL196, NGL049, NGL057, NGL216, ICBG712 and ICBG721) harbor more BGCs than more derived strains. Within the *Escovopsis* spp. there is an increase in BGC abundance from the most basal strains (coral A) to the more derived (leaf cutter agriculture) (Figure S4C).

**Figure 3.**
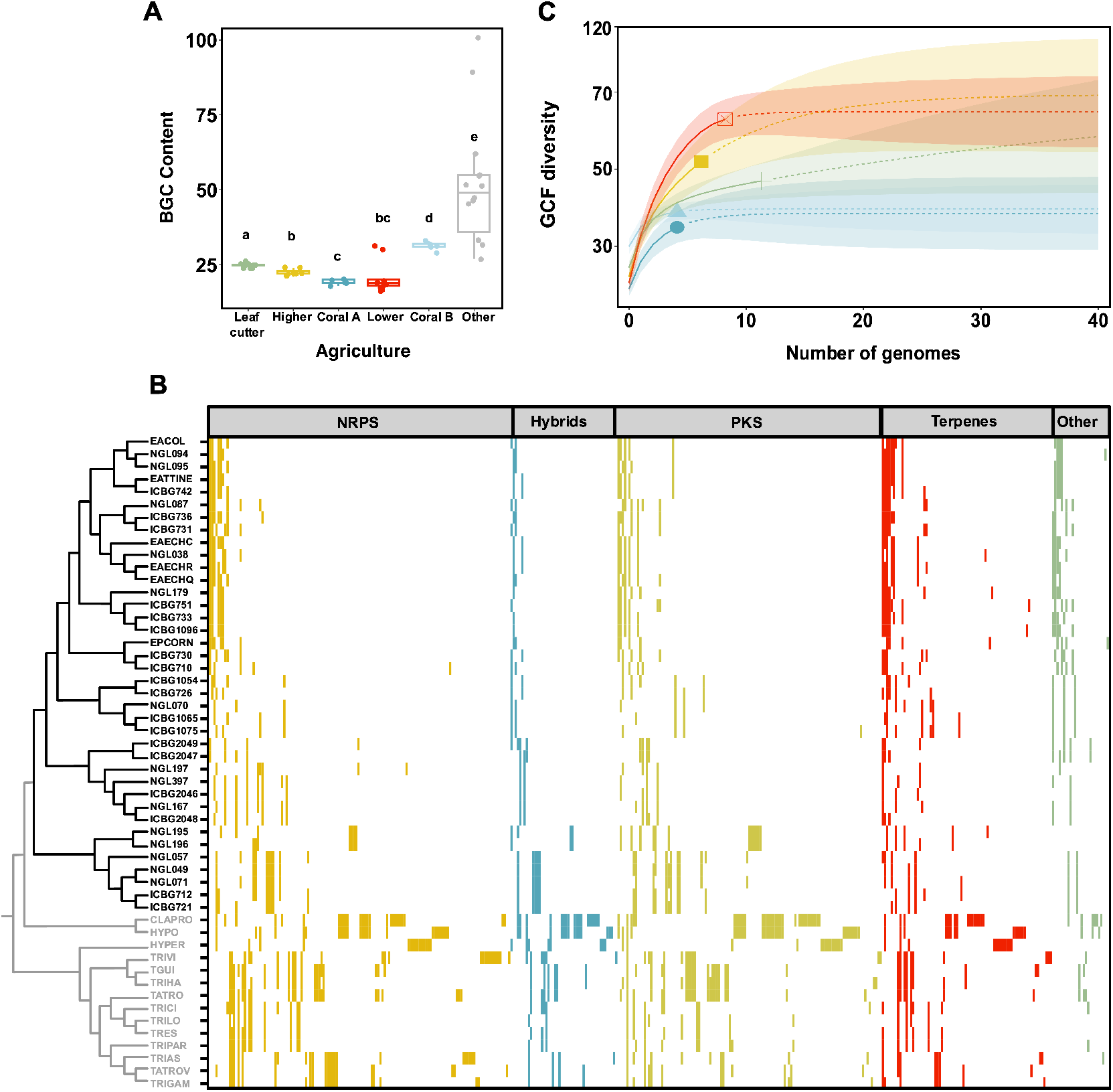
(A) Total number of biosynthetic gene clusters (BGCs) identified across escovopsis strains infecting different attine agricultural systems and non-ant associated relatives. Box colors denote attine clades systems: grey, free-living; green, leaf-cutter agriculture (*Escovopsis* spp.); yellow, general higher agriculture (*Escovopsis* spp.); blue, coral agriculture A (*Escovopsis* spp.); red, lower agriculture (*Sympodiorosea* spp.); and light blue, coral agriculture B (undescribed genus). Graph background is colored according to the general agriculture strains exploit: yellow, higher agriculture; and red lower agriculture. (B) Gene Cluster Family (GCF) distribution across escovopsis and closely related fungi. Each column in the heatmap represents a GCF. The presence of a GCF in a strain is highlighted by colored blocks according to BGC class: yellow, NRPs blue, PKS-NRPS hybrids; light green, PKS; red, Terpenes; and green, others (including RiPPs, indoles, siderophores and others). The absence of a GCF is represented by white spaces. (C) Rarefaction curves assessing GCF richness in escovopsis strains across different attine agricultural systems for the given sequencing effort. Continuous lines represent observed diversity, dashed lines inferred diversity. Shaded areas denote confidence intervals. Colors denote agricultural systems: green, leaf-cutter agriculture, yellow, general higher agriculture; blue, coral agriculture A red, lower agriculture; and light blue, coral agriculture B.

The reduction in BGC abundance in escovopsis relative to other non-ant associated Hypocreales is consistent with its shift in lifestyle to an obligate symbiont of ant nests. Transitions from free-living states to obligate symbioses can often be accompanied by gene loss due to relaxed selection on genes that are no longer necessary in a stable, predictable environment (24, 92). Additionally, some specialist parasites are known to harbor a narrower suit of BGCs relative to generalist ones. For instance, *Metarhizium* strains that acquired the *dtx* biosynthetic gene cluster, responsible for the synthesis of a diversity of toxins, have broader host ranges (infecting hundreds of insect species) compared with non-toxigenic strains (lacking the BGC) that present much narrower host ranges, infecting only locusts and grasshoppers (4). Correlating with a higher content in biosynthetic gene clusters, *Escovopsis* spp. strains infecting higher agriculture (e.g. *E. weberi*) are thought to be more virulent than escovopsis infecting lower agriculture (19).

To compare BGC composition across strains, we grouped BGCs into gene cluster families (GCFs) based on sequence homology and cluster architecture employing the BiG-SCAPE algorithm. The resulting sequence similarity network built with a similarity score cutoff of 0.5, clustered 1595 BGCs into 411 gene cluster families (GCFs). We visualized the GCF distribution across escovopsis through the construction of a presence/absence table (Figure 3B). One hundred and twenty-eight GCFs were present in the sampled escovopsis, and 102 of them were unique to escovopsis relative to non-Attine associated fungi. Only 26 GCFs were shared between escovopsis and other Hypocreales species (Table S8, Figure 3B). A rank-abundance curve demonstrates that 27 GCFs occur only once in escovopsis, and an additional 27 are present in just two strains (Figure S5). Surprisingly, no GCF as defined by BigSCAPE was ubiquitous in escovopsis, and therefore characteristic of the group of symbionts as a whole. Rarefaction curves provide an assessment of GCFs richness for the given sequencing effort and reveal that although our sampling was largely adequate, additional chemical diversity is yet to be discovered, especially within the most basal clade of escovopsis infecting lower attine gardens (Figure 3C). Further sequencing efforts in strains from this group will surely reveal additional GCFs.

To distinguish novel BGCs from already described ones, we supplemented our dataset with characterized gene clusters from the MIBiG database as a reference, which at the date of publication contained 1923 BGCs, out of which 207 were of fungal origin. Only five GCFs in our escovopsis dataset are homologous to BGCs in the database. Three families comprising highly similar BGCs cluster together with the MIBiG cluster BGC0001585, responsible for the synthesis of melinacidin IV, suggesting they represent slightly different variants of the same biosynthetic pathway. The other two GCFs are homologous to BGC0001583 and BGC0001777, which encode for emodin and shearinines respectively. The distribution of all three GCFs is discrete. While most strains of escovopsis harbor the BGC responsible for the production of melinacidin IV, those encoding for shearinine and emodin are restricted to more derived clades (i.e., *Escovopsis* spp. for shearinine, and *Escovopsis* spp, with the exception of those exploiting coral agriculture for emodin).

Fermentation experiments using *E. weberi* have led to the detection and potential functional role of all three metabolites and some derivatives (26). *E. weberi*-produced shearinine derivatives can deter ants and are lethal at high concentrations, preventing insect workers from weeding their garden, thus allowing the parasite to persist in the nest (26). The production of the epipolythiodiketopiperazine melinacidin IV inhibits the growth of the ant defensive mutualist *Pseudonocardia*, whereas the synthesis of emodin has detrimental effects on the cultivar (26) and other co-occurring actinobacteria, such as *Streptomyces*. While the production of these metabolites has been detected in *Escovopsis* strains parasitizing leaf cutter ant gardens, our results demonstrate that the distribution of these BGCs is broader than previously thought and extends to strains exploiting other agricultural systems. Whereas shared GCFs with other fungal genera suggests that they may play a general role in fungal physiology, the presence of GCFs characteristic of specific clades correlating with different attine agricultural systems likely reflects the distinct selective pressures exerted on the symbionts by these different ecosystems. These results are consistent with an on-going arms-race where these symbionts must constantly evolve new adaptations to overcome not only cultivar defenses, but also, very likely, ant defenses, those exerted by protective symbionts such as *Pseudonocardia* and those exerted by other microbes that inhabit these complex microbial communities. For example, the defensive symbiont of beewolves, *Streptomyces* spp., produces different antibiotic cocktails (both in composition and concentration) in association with each insect species, but also in distinct geographical regions (93), presumably as an adaptation to defend their hosts against different local pathogen communities. Furthermore, the variation metabolic profiles of escovopsis could be a a reflection of them having different impacts on the symbiosis; while some (e.g., *E. weberi*) have been shown to be highly virulent parasites of the ants’ cultivars, experimental tests of the impacts of most species are not known. More experimental work is required to assess the specific roles that individual metabolites may play in the ecology of this diverse group of symbionts..

Recent surveys reveal that less than 3% of the biosynthetic space represented by fungal genomes has been linked to metabolites (91, 94). As such, most of the GCFs identified by BGC similarity networks could not be correlated with known compounds, suggesting *Escovopsis* is a promising source for the discovery of new bioactive compounds.

### Gene cluster families (GCFs) delineate groups of *Escovopsis*

To assess differences in biosynthetic profiles between *Escovopsis* strains exploiting different attine agricultural systems we performed a non-metric multidimensional scaling analysis. Our results demonstrates that *Escovopsis* strains harbor very different GCF profiles than related non-ant associated fungi, and that these profiles differ between *Escovopsis* infecting higher and lower agriculture (Figure 4A, ANOSIM, R = 0.59, p < 0.001, 999 permutations). Likewise, GCF profiles are sufficient to cluster strains into separate groups based on phylogenetic lineage (Figure 4B, ANOSIM, R = 0.87, p < 0.001, 999 permutations). A PERMANOVA analysis reveals that most of the variation (95%) is explained by the interaction between symbiont genus and ant species (Figure 4B, adonis2, 999 permutations, R^2^=0.952, p=0.001).

**Figure 4.**
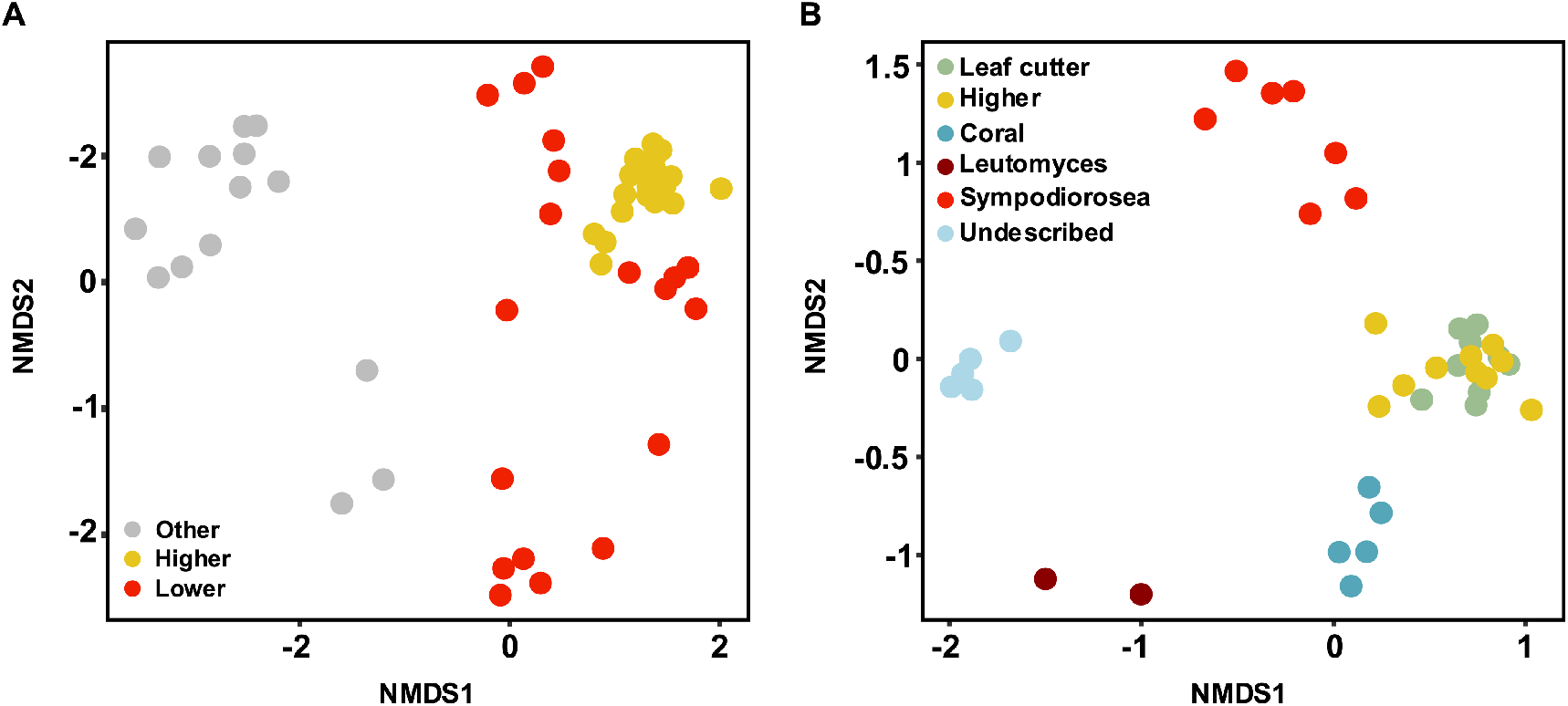
Gene cluster families (GCFs) delineate groups of escovopsis strains. (A) Non-metric multidimensional scaling (NMDS) plot showing differences in GCF composition among escovopsis infecting higher agriculture (yellow) and lower agriculture (red) and non-ant associated fungal relatives (gray). (B) NMDS plot depicting GCF composition in escovopsis strains across ant clades: *Escovopsis* spp. (green, leaf-cutter agriculture, yellow, general higher agriculture; blue, coral agriculture A), dark red, *Luteomyces*; red, *Sympodiorosea*; and light blue, the undescribed genus.

Based on the presence/absence matrix of GCFs across strains, we constructed a hierarchical clustering analysis. *Escovopsis*’ phylogeny, based on all orthologs, and the GCF dendrogram are highly congruent (Figure 5A), with the exception that the most basal clade of strains parasitizing coral agriculture and lower agriculture appear as paraphyletic in the GCF dendrogram. An entanglement analysis, which gives a visual approximation of the level of agreement between two dendrograms (95), yielded a score of 0.02, suggesting a high degree of congruence between escovopsis*’* phylogeny and the GCF dendrogram.

**Figure 5.**
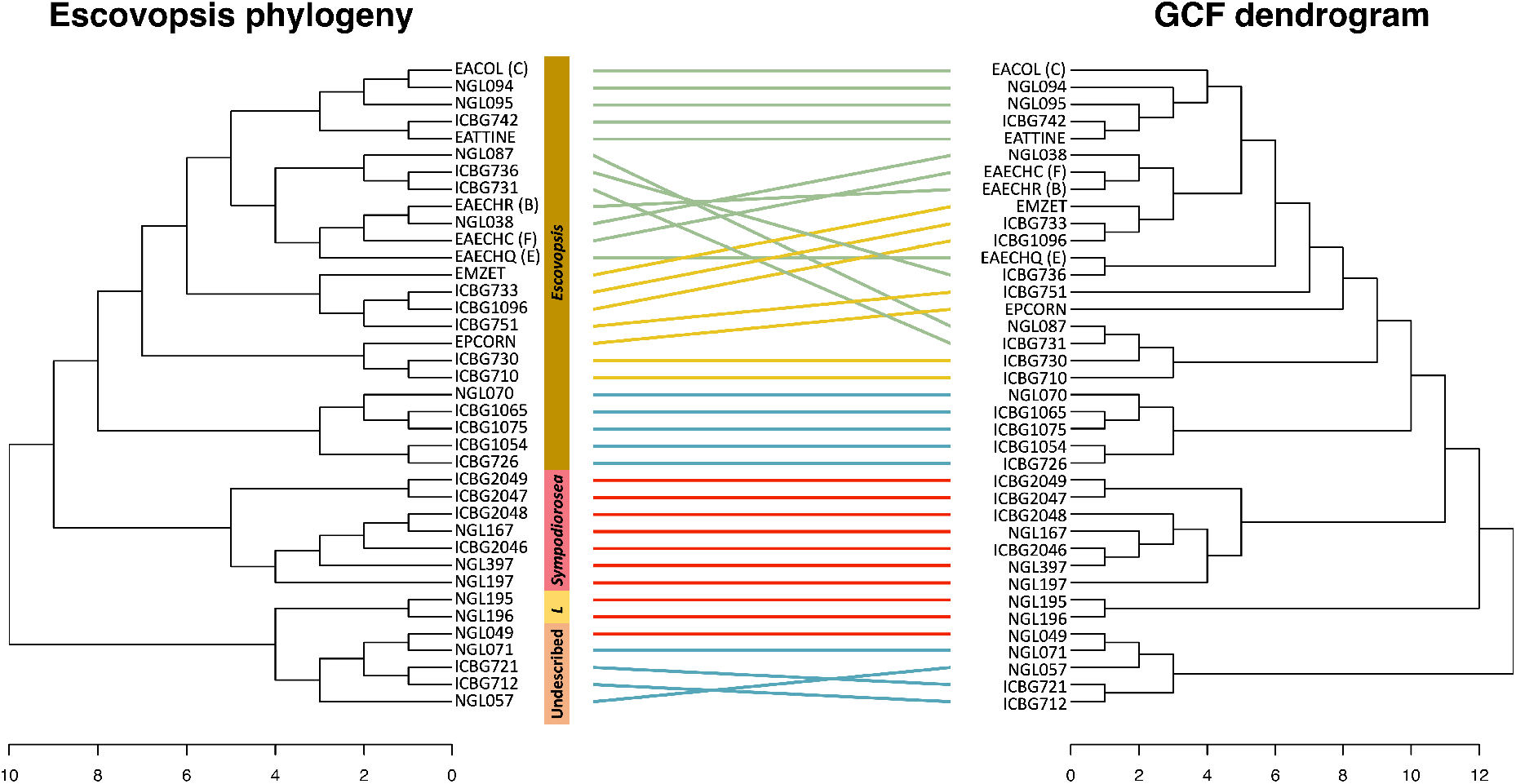
GCF profiles in *Escovopsis* are a phylogenetic trait. Tanglegrams revealing congruence between (A) *Escovopsis* phylogeny and strain biosynthetic potential. Lines connect strains with their GCF profile and colors denote attine clades systems: grey, free-living; green, leaf-cutter agriculture, yellow, general higher agriculture; blue, coral agriculture A red, lower agriculture; and light blue, coral agriculture B. Branches have been rotated for maximum congruency. The maximum likelihood tree was built with 681 single copy orthologous genes. The chemical dissimilarity dendrogram was generated using hierarchical cluster analysis on the presence and absence of GCFs using Jaccard distance and UPGMA (unweighted pair group method with arithmetic mean) as the agglomeration method.

These results suggest that the parasite’s biosynthetic potential is a phylogenetic trait and can be employed to delineate groups of escovopsis, particularly at broad taxonomical levels. Christopher et al. (69) demonstrated that phylogenetic analyses based on chemical profiles of *Escovopsis*, resulted in similar tree topologies to gene-based phylogenies, confirming that chemical profiles can be considered phylogenetic traits. Additionally, the congruency between the species phylogeny and the BGC profile dendrogram, suggest that BGCs are evolving in parallel with escovopsis species, and that pathway gains and subsequent vertical inheritance, as well as losses are the main forces driving BGC diversification, given that common horizontal transfer of BGCs between strains would result in incongruent topologies.

### a. Pathway evolution: Ancestral state reconstruction

To explore the evolutionary history of escovopsis’ biosynthetic pathways relative to their encoding strains, we performed an ancestral state reconstruction analysis. We clustered GCFs into pathways (Ps) based on the assumption that they produce related compounds (See methods section, Table S9). The 411 GCFs detected in our escovopsis dataset were clustered into many different pathways. Their distribution was overlaid onto a simplified parasite phylogeny, generated by collapsing certain branches on the species tree, resulting in 8 lineages A-H, that correspond with the newly described genera (A, undescribed genera; B, *Luteomyces*; C, *Sympodiorosea*; and D-H *Escovopsis* spp.) Figure 6). 67 pathways were present in escovopsis, out of which 56 were unique to this group of symbionts and 11 were shared with other hypocreales. The analysis revealed that 15 pathways were present in the common ancestor of escovopsis, and 11 of those were shared with the closely related genus *Cladobotryum.* The transition from a non-ant associated lifestyle to a fungal garden parasite correlates with the loss of one pathway (P67), involved in the biosynthesis of an uncharacterized PKS, that is present in all close relatives but absent in every *Escovopsis* strain. Five pathways (P7, P8, P10, P14 and P15) evolved early in the evolutionary history of these fungal symbionts and are present in most strains. However, none of them are ubiquitous, as there have been some clade-specific loses.

**Figure 6.**
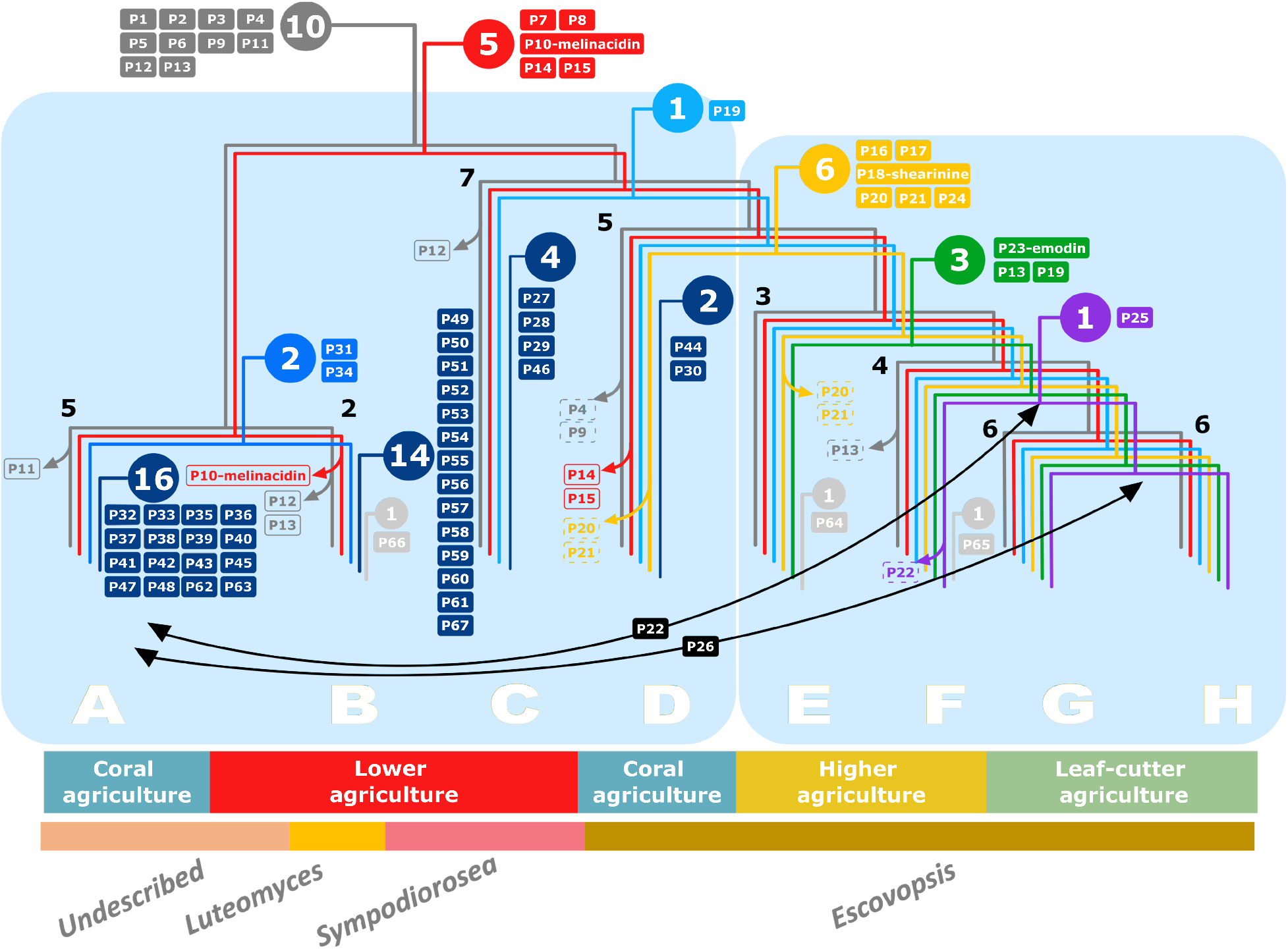
Phylogeny depicting the inferred ancestry of secondary metabolite biosynthetic pathways present in escovopsis. A simplified species phylogenomic tree depicts 8 major *Escovopsis* lineages A-H, that correspond to the newly proposed taxonomical divisions. The number of the strains in each group is indicated in black adjacent to branch nodes. Circles indicate the number of pathways originated at various points in the species tree, whereas filled boxes state pathways next to the point of acquisition. Transparent boxes represent pathway losses in all strains within a clade (continuous outline) or more than 50% of the strains in the clade (dashed outline). Grey, pathways shared with the sister clade; red, shared the common ancestor of the genus; indigo, present in basal strains; light blue, present in derived clade; yellow; shared between strains infecting derived coral agriculture and general higher agriculture; green, shared by all higher agriculture; purple, shared by derived general higher agriculture and leaf-cutter agriculture; dark blue, clade-specific pathways; light gray, strain-specific pathways. Black arrows describe HGT events.

The remaining 50 pathways were acquired at various points during the evolution of the genus either through horizontal gene transfer (HGT) or *de novo*. An average of three pathways are acquired by *Escovopsis* with every transition to a new ant agricultural system. However, the transition from strains within the three most ancestral escovopsis clades (A-C, *Sympodiorosea*, *Leutomyces* and an undescribed genus) to the most derived super-clade including clades D-H (*Escovopsis* spp.) correlate with the acquisition of 5 pathways, including P18, responsible for the biosynthesis of shearinine D and indicates that these pathways are unique for *Escovopsis* spp. Four pathways, including P24 which synthesizes emodin, evolved early in the divergence of *Escovopsis* to infect higher attine agriculture. Interestingly, no pathway is unique to the most derived clade of the parasite (i.e., strains parasitizing leaf-cutter ant nests, clades G and H).

Phylogenetic analysis of key biosynthetic genes from each pathway confirms, based on congruence with the species tree, vertical inheritance for most of the pathways following acquisition. However, they also suggest that some pathways were exchanged between strains. P22, encoding for a terpenoid, has been transferred between the ancestor of parasite strains exploiting higher attines (ancestor of clades F-H) and the basal clade parasitizing coral agriculture (clade A). Similarly, P26, encoding a PKS, has been shared between the ancestor of strains infecting leaf-cutter agriculture and the most derived clade parasitizing coral agriculture. In both cases, the direction of the exchange remains unclear. However, once transferred, these pathways have subsequently been vertically inherited by all members of the clades.

The evolution of *Escovopsis’* biosynthetic potential has not only evolved through pathway acquisition, but also through BGC loses. Six pathways have been lost in strains infecting lower attines: three that were already present in the sister clade represented by *Cladobotryum* and *Hypomyces rosellus* (P10, 11, 12, and P13) and three that evolved in the common ancestor of all escovopsis strains (P10, P14 and P15). P12 appears to have been lost twice, once in clade B (*Luteomyces*), as well as in clade C (*Sympodiorosea*). P4 and P9 have also been lost in 4 and 2 strains respectively. Within *Escovopsis* parasitizing higher attines, no pathway has been lost completely. Only three pathways have been lost in some strains: the ancient P13 in clade F, and the more recently evolved P18, encoding for shearinine, in one clade E strain (EPCORN). While the loss of this BGC in EPCORN, and its inability to synthesize the resulting compound was already described through both bioinformatic and chemical assays (26), our results suggest it is not a widespread event, given that all the remaining strains still conserve the BGC. A number of pathways (P4, P9, P14, P15, P20, P21) have been lost in *Escovopsis* spp. strains that experienced a host-shift, from exploiting a *Leucocoprineae* to a *Pterulaceae* host. It is plausible that these pathway losses represent an adaptation and specialization to exploit a new host. In general, more pathways have been lost in *Escovopsis* strains infecting lower attine gardens than those attacking the cultivars of higher attines, and those pathways were most often ancient, suggesting that newly acquired BGCs either (i) have not had enough evolutionary time to be selected against, or (ii) may be adaptive and thus maintained in *Escovopsis*. These results oppose patterns described in other fungi, where generalist parasites harbor more BGCs than specialist ones (4). Escovopsis strains attacking lower attines are thought to be less specialized than those infecting higher attine gardens (19). However, our results suggest that they may be more specialized than previously thought. Furthermore, the colonies of lower attines are smaller than those of higher attines, consisting of a handful and millions of workers respectively. Given the insecticidal properties of some BGCs, it is plausible that parasite strains attacking bigger colonies, require a more diverse cocktail of bioactive compounds relative to those infecting smaller colonies. In fact, studies have demonstrated that the proportion of ant nests harboring fungal contaminants (fungi other than the cultivar) is highest in lower attines (13). However, the proportion of those contaminants made up by *Escovopsis* and its allies is highest for higher attines (13). This could be the result of *Escovopsis* attacking higher attine nests being able to better fend off both fungal competitors than those infection lower attine nests, and inhibiting ant-weeding behavior, given their higher content in BGCs. Additionally, *Escovopsis* strains infecting small colonies may encounter less diverse microbial communities in compared to those encountered in bigger gardens, and may not require as many antibiotic compounds to outcompete other microbes.

Our results suggest that *Escovopsis* acquired the capability to synthesize the antimicrobial compound melinacidin IV early in the evolution of this group of symbionts and was subsequently lost in lineage B (P10), i.e. *Luteomyces* parasitizing lower attines. The evolution of the pathway, is however, uncertain. Although we did not detect the presence of the core biosynthetic enzymes in the sister clade to escovopsis, consisting of *Cladobotryum* and *Hypomyces* strains, other Hypocreales are known to synthesize this metabolite, such as *Acrostalagmus* sp, a rare fungal genus that has been found associated with soil (96), mushrooms (97) and plant material (98), suggesting this BGC may have been acquired horizontally. However, while the closely related genus *Trichoderma*, has never been described to synthesize this antibiotic, strains within this genus harbor a number of homologous genes to the melinacidin IV BGC, including the backbone enzyme (99). Therefore, it is alternatively plausible that the pathway responsible for the production of melinacidin IV evolved early within the hypocreal family and was lost in the *Cladobotryum-H. rosellus* clade, accumulating enough changes (or requiring fewer genes than previously thought) that we have classified them as different GCFs in our survey.

The inferred ancestry for the pathway responsible for shearinine (P18) biosynthesis, suggests it was characteristic of *Escovopsis* spp. While absent from other Hypocreales, a BGC encoding for shearinine D, has been described for the distantly related fungus *Penicillium janthinellum* (100), suggesting that it may evolved through HGT in these symbionts. Emodin, encoded by pathway P24, was one of the last BGCs to evolve within *Escovopsis*, appearing in the ancestor of strains parasitizing general higher agriculture and leaf-cutter ants.

The evolutionary transition between lower to higher agriculture in attine ants correlates not only with an increment colony size (from hundreds to millions of workers) (84), but also with an incipient division of labor between worker ants that culminates with the cast system in leaf-cutter ants (101). The transition of escovopsis from attacking lower to higher attine agriculture coincided with the evolution of a new suit of biosynthetic gene clusters, possibly explaining the increase in complexity required by escovopsis to survive in this environment, and the split of these strains into different genera.

## 4. Conclusion

Parasites evade and counter host defenses through a remarkable array of secondary metabolites and natural products. Here, we highlight the highly streamlined genomic features of escovopsis, defined by 7 chromosomes, harboring few repetitive sequences. Despite a high degree of metabolic conservation, we observe limited variation in the parasite’s potential to produce secondary metabolites. As the phylogenetic distribution of the encoding biosynthetic gene clusters coincides with attine transitions in agricultural systems and cultivar types, we highlight the likely role of these metabolites in mediating adaptation by a highly specialized pathogen. Future efforts will shed light on the mode-of-action and mechanistic basis of *Escovopsis* secondary metabolites, their role in underlying virulence and pathogenicity in an ancient agricultural system.

## 5. Acknowledgements

We acknowledge funding from the German Research Foundation (Deutsche Forschungsgemeinschaft) under the individual grant program (BE6922/1-1) and Germany’s Excellence Strategy (EXC 2124 – 390838134) for AB. This work was also funded by the National Science Foundation (award 1711545) for CC, (award 1754595) for HS, NMG and TR, and (award1927411) for NMG and AR, QVM and AR would like to thank the São Paulo Research Foundation (Fundação de Amparo à Pesquisa do Estado de São Paulo [FAPESP], grant number 2019/03746-0. We are very thankful to Dr. Martina Adamek for advice on BGC clustering and Dr. J Lovell and Dr. Navarro for help in troubleshooting software functioning.

## 6. Data accessibility

Accession numbers will be provided upon acceptance. BigScape results and other supplementary material can be found in FigShare under the title’s project.

## 7. Author contributions

AB, HS, NMG and NZ conceived of the study. NMG and YC collected samples. AB, HS and CC performed DNA extractions. AB, HS and AGM assembled genomes. AB, QV and AR performed analyses. AB wrote the manuscript. All authors provided valuable comments to the manuscript.

